# FAM210B Regulates Iron Homeostasis and Sex-Specific Responses in Stress Erythropoiesis

**DOI:** 10.1101/2023.09.26.559581

**Authors:** Mark Perfetto, Muhammad Ishfaq, Aiden Mohideen, Catherine M. Rondelli, Samantha Gillis, Jesus Tejero, Amber N. Stratman, Rebecca Riggins, Yvette Y. Yien

## Abstract

Iron is required for redox homeostasis but poses toxicity risks due to its redox activity. Erythropoiesis hence requires tight regulation of iron utilization for hemoglobin synthesis. The requirement for iron in erythropoiesis has necessitated the evolution of mechanisms to handle the iron required for hemoglobinization. FAM210B was identified as a regulator of mitochondrial iron import and heme synthesis in erythroid cell culture and zebrafish models. Here, we demonstrate that while FAM210B is required for erythroid differentiation and heme synthesis under standard cell culture conditions, holotransferrin supplementation was sufficient to chemically complement the iron-deficient phenotype. To investigate the role of FAM210B in erythropoiesis, we used knockout mice. While *Fam210b^−/−^* mice were viable and did not exhibit overt erythropoietic defects in the bone marrow, the male mice exhibited an increase in serum transferrin suggesting sex-specific alterations in systemic iron sensing. Upon phlebotomy-induced stress erythropoiesis, *Fam210b^−/−^* mice exhibited differences in serum transferrin levels, and more starkly, had markedly smaller spleens indicating defects in stress response. *Fam210b^−/−^* males had defects in neutrophil and monocyte numbers, as well as decreased erythroid progenitor numbers during erythropoietic stress. Together, our findings show that *Fam210b* plays a key role in splenic response to erythropoietic stress Our findings reveal a critical role for FAM210B in mediating splenic stress erythropoiesis and suggest it may act as a sex-specific regulator potentially linked to androgen signaling.

## Introduction

Iron is an essential redox element in iron-sulfur clusters, heme, and iron-binding proteins (1,2). Iron is of especial importance in erythroid biology, most prominently for hemoglobin production, but also regulating processes related to terminal erythropoiesis such as proliferation and nuclear condensation (3–5). Iron-containing proteins are required for numerous other cell-specific processes such as dopamine production (6,7), adipocyte function (8,9) and proliferation of hepatocytes in response to injury (10), as well as essential “housekeeping” processes such as mitochondrial respiration and DNA synthesis (11,12). Despite the the requirement of iron in diverse cellular processes, the mechanisms governing its distribution across tissues and its intracellular fate remain poorly understood.

*Fam210b* was previously identified to be a transcriptional target of EPO and GATA1 whose expression was enriched in erythroid cells (5,13). Erythroid FAM210B is an inner mitochondrial membrane protein that facilitates the mitochondrial import of iron and its incorporation into heme (5,13). FAM210B regulates metabolism in several tissues (5,13-15) and its biology is complex. In ovarian cancer cells, FAM210B promotes glycolysis (14). In *C. elegans*, a FAM210 homolog is required for reproduction via its regulation of oogenesis (16). More recently, Suzuki et al. demonstrated that when HiDEP cells--fetal globin expressing erythroid cells derived from induced pluripotent cells (iPS)--are differentiated in the presence of holotransferrin and 20% FBS, the absence of FAM210B actually *promotes* erythroid terminal differentiation (15). These divergent findings suggested that FAM210B function is context specific and dependent on tissue type, source of tissue, and nutrient conditions.

Currently published studies on vertebrate FAM210B have largely been performed on cells in culture. We previously determined that FAM210B was required for maximal heme synthesis, proliferation, and maintenance of cell number in *Fam210b* knockdown primary murine fetal liver erythroid cells in which and zebrafish embryos (5). While we completed this work to describe the role of FAM210B in fetal/embryonic tissue, no work had addressed the role of FAM210B in mammals. To further investigate the *in vivo* function of FAM210B, we used the whole body *C57BL6N/6N-^Atm1Brd^Fam210b^tm1b(KOMP)/Mbp^* /MbpMmucd mouse line established by the Mutant Mouse Resource and Research Center (MMRRC) at University of California at Davis. Our results indicate that at steady state, FAM210B is dispensable for erythroid differentiation and heme synthesis in the adult bone marrow under normal conditions. *Fam210b^−/−^* mice are viable and reproduce normally, unlike their *C. elegans* counterparts. At steady state, their hematopoietic indices, including hemoglobin and erythrocyte count, were within normal parameters, although we observed significantly increased serum transferrin in the males, suggesting that they were responding to an iron deficiency signal. Given our previously published observations that FAM210B deficiency in cell culture and zebrafish embryos caused anemia, we proceeded to determine if FAM210B played a role in erythropoiesis during iron deficiency or erythropoietic stress. Although red cell count, hemoglobin levels and bone marrow erythropoiesis did not differ between phlebotomized WT and *Fam210b^−/−^* mice, phlebotomized *Fam210b^−/−^* mice had decreased spleen size relative to their body weight suggesting a defect in erythropoietic recovery from stress. In particular, male *Fam210b^−/−^* mice had decreased numbers of splenic erythroid progenitors, but increased numbers of splenic reticulocytes relative to wild-type, suggesting that terminal erythroid differentiation was accelerated at the expense of maintenance of progenitor population pools, potentially as a mechanism to maintain erythroid cell numbers during stress. These findings were consistent with previously published data in human erythroid HiDEP cells derived from induced pluripotent cells (15). Although *Fam210b^−/−^* females also had decreased spleen weights, we did not observe similar changes in erythroid progenitor proportions. Collectively, our data indicates that in adult mice, FAM210B plays a male-specific role in iron homeostasis and recovery from erythropoietic stress, and additionally plays a previously unidentified role in non-erythroid autonomous splenic response to erythropoietic stress. These findings reveal the first known mechanism of iron and erythroid homeostasis that is both sex-specific and stress-responsive and highlighting the critical importance of investigating physiological phenomena in both sexes.

## Results

To determine the extent to which immortalized erythroid cells model the role of FAM210B in erythropoiesis, we verified that differentiated *Fam210b^−/−^* murine erythroleukemia (MEL) cells are hemoglobin deficient (Figure 1A, B). MEL cells are Friend virus transformed, spleen derived erythroleukemia cells that can be induced to differentiate by addition of polar compounds like DMSO (17). Both non-differentiated and differentiated *Fam210b^−/−^* MEL cells are larger than wild-type cells, indicating a defect in erythroid terminal differentiation (Figure 1C). We previously described FAM210B’s main erythroid phenotype as a defect in mitochondrial iron metabolism under cell culture conditions where the only iron sources were DMEM and 10% FBS (5). Suzuki *et al.* reported that HiDEP erythroid cells were differentiated in serum containing 20% FBS and a transferrin supplement, which provides significantly more bioavailable iron than what was used in our previous work (15). Therefore, we postulated that increased iron content in HiDEP cell differentiation media could have chemically complemented the iron deficiency and hemoglobinization defects in *Fam210b^−/−^* cells, masking phenotypes we had previously observed (5).

**Figure 1.**
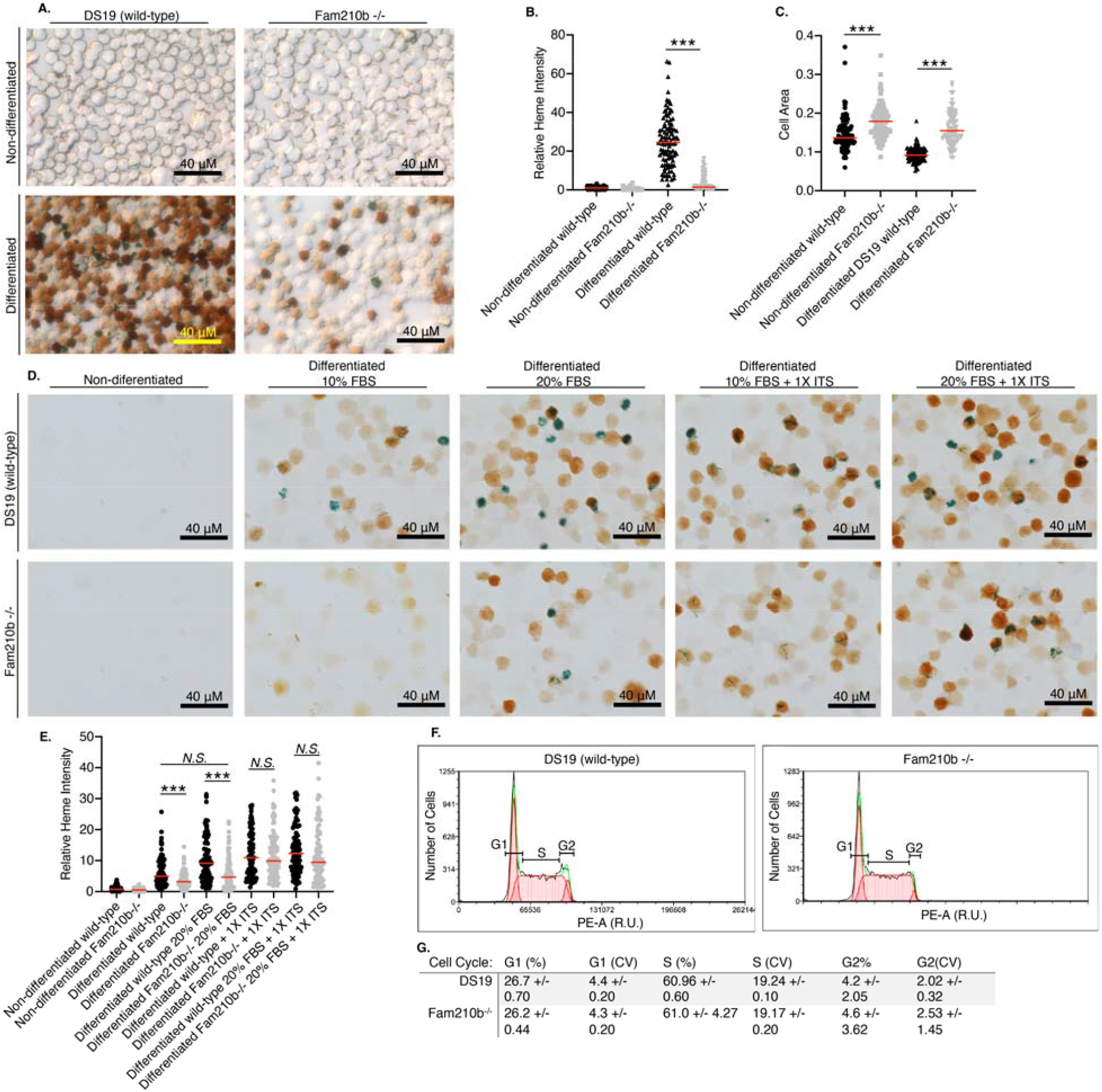
Fam210b is required for heme synthesis and normal cell size in MEL cells. **A.**, 63X brightfield image of non-differentiated and differentiated MEL cells stained for heme (brown color), N=3 from 3 experiments.**B.**, Relative heme stain intensity of the MEL cells from A. Each point represents one cell, n=102 cells from 3 separate biological replicates. **C.**, Cell areas of Fam210b^−/−^ MEL cells were significantly larger than WT cells. **D.** 63X brightfield image of non-differentiated and differentiated MEL cells incubated with fetal bovine serum (FBS) and insulin-transferrin-selenium (ITS) and stained for heme (brown/green color). **E.**, Relative heme intensity of MEL cells from D (n=100). **F.**, Cell cycle analysis of WT and Fam210b^−/−^ MEL cells, N=3. **G.**, Quantitation of cell cycle stage progression. **\*\*\***, p<0.001; N.S., not significant.

To replicate the findings of Suzuki *et al.*, we cultured *Fam210b^−/−^* MEL cells in 10% FBS with transferrin supplement, 20% FBS alone, or 20% FBS with transferrin supplement and quantitated benzidine staining as a metric of hemoglobinization. We decreased the staining time to 5 min to allow us to observe increases in hemoglobin content caused by iron treatment (Figure 1D, E). Consistent with our previous results, there was a significant decrease in hemoglobinization in *Fam210b^−/−^* cells relative to wild-type cells when grown in either 10% or 20% FBS. However, addition of a holotransferrin supplement increased the hemoglobinization in *Fam210b^−/−^* cells such that their benzidine staining signal does not significantly differ from wild-type cells. These data indicate that the elevated levels of transferrin-bound iron in HiDEP cell culture media may have masked phenotypes that manifest during differentiation under iron-deficient conditions, where FAM210B’s role is likely to be most important. While the work on *Fam210b* deficient HiDEP cells does support that FAM210B is required for mitochondrial function and metabolism, it is interesting to note that other mitochondrial respiratory complex proteins were upregulated in *Fam210b* deficient HiDEP cells, suggesting a level of genetic compensation is occurring within these cells to allow for erythroid cell survival and differentiation. Finally, because *Fam210b^−/−^* primary erythroid cells experienced proliferation defects, we were curious to determine if this defect could be recapitulated in *Fam210b^−/−^* MEL cells. MEL cells did not recapitulate the cell cycle defects in primary cells, indicating the limits of immortalized cell lines as experimental models (Figure 1F). Therefore the finding holotransferrin is sufficient to rescue hemoglobinization in *Fam210b^−/−^* cells to wild-type levels is particularly important to note when studying erythroid cells, which predominantly rely on transferrin bound iron for hemoglobin synthesis (18).

There are few published studies on the *in vivo* role of FAM210B. We previously attempted to generate zebrafish *fam210b* knockout mutants using CRISPR/Cas9 with no success. The genomic region was poorly annotated and contains extensive repetitive regions, making gRNA design and genotyping challenging. Our zebrafish experiments therefore used morpholino-mediated gene suppression (5) to investigate the role of *Fam210b* in zebrafish development. We showed that *fam210b* was required for embryonic erythropoiesis and erythroid heme synthesis and that iron supplementation could restore erythropoiesis in *fam210b*-deficient zebrafish (5). However, we were not able to follow the full course of development to adulthood because of the transient nature of morpholino-mediated knockdown.

To address this gap, we used *C57BL6N/6N-^Atm1Brd^ Fam210b^tm1b(KOMP)/Mbp^*/MbpMmucd (from here on *Fam210b^−/−^*) knockout mice generated by the Mutant Mouse Resource and Research Center (MMRRC) at the University of California at Davis. Crosses between *Fam210b^+/−^* mice resulted in 25% of pups with the *Fam210b^−/−^* genotype, indicating they were viable without noticeable divergence from the expected mendelian ratio. To determine if FAM210B was required for fertility, we crossed *Fam210b^+/−^* to *Fam210b^−/−^* mice which resulted in 48.4% *Fam210b^−/−^* offspring, indicating that *Fam210b^−/−^* did not have fertility defects, that *Fam210b*^−/−^ mice did not have fertility defects, in contrast with the observed differences by Kang et al. in C elegans, where it FAM210 is required for oogenesis and reproduction (16). Zebrafish *Fam210b* is maternally expressed (5). To determine if maternal *Fam210b* was required for the survival of *Fam210b^−/−^* mice, we bred *Fam210b^−/−^* mice to one another. We obtained 100% of *Fam210b^−/−^* pups, indicating that maternal FAM210B is not required for survival of *Fam210b^−/−^* embryos (Table 1). Since *in vivo* erythropoiesis in *Fam210b^−/−^* fetuses was sufficient for their survival, we did not conduct studies on fetal liver erythropoiesis. *Fam210b^−/−^* mice survived until they were euthanized at about 1 year of age.

**Table 1.**
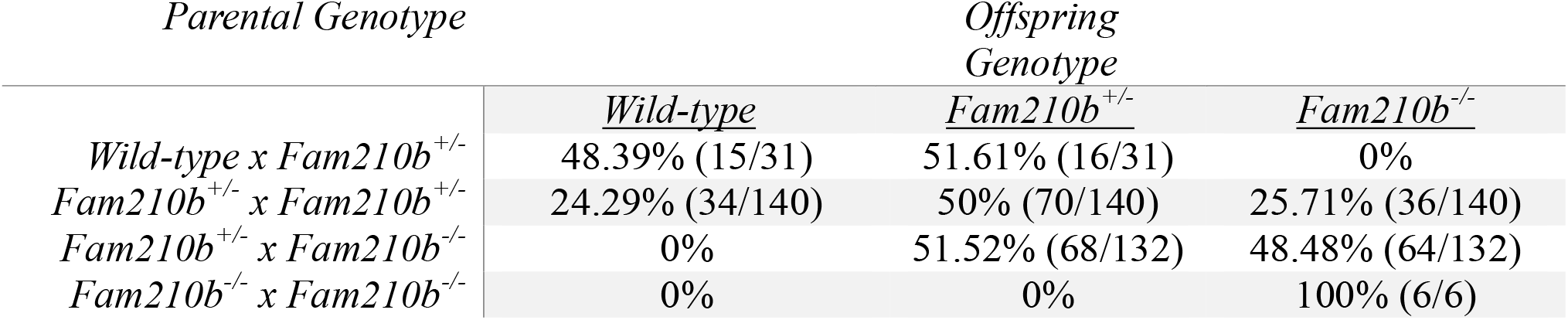
Proportions of offspring genotypes obtained from matings of *Fam210b^+/−^* or *Fam210b^+/−^* mice.

| *Parental Genotype* | *Offspring Genotype* |  |  |
| --- | --- | --- | --- |
|  | <u>*Wild-type*</u> | <u>*Fam210b*<sup>+/-</sup></u> | <u>*Fam210b*<sup>-/-</sup></u> |
| *Wild-type* x *Fam210b*<sup>+/-</sup> | 48.39% (15/31) | 51.61% (16/31) | 0% |
| *Fam210b*<sup>+/-</sup> x *Fam210b*<sup>+/-</sup> | 24.29% (34/140) | 50% (70/140) | 25.71% (36/140) |
| *Fam210b*<sup>+/-</sup> x *Fam210b*<sup>-/-</sup> | 0% | 51.52% (68/132) | 48.48% (64/132) |
| *Fam210b*<sup>-/-</sup> x *Fam210b*<sup>-/-</sup> | 0% | 0% | 100% (6/6) |

We verified knockout of *Fam210b* expression in mice by semi-quantitative RT-PCR of *LacZ* (Figure 2A) and suppression of the *Fam210b* transcript using qRT-PCR (Figure 2B). We also verified knock-out of *Fam210b* expression by PCR for *Fam210b* from genomic DNA extracted from ear clips (Figure 2C). Male *Fam210b^−/−^* mice were heavier than wild-type and heterozygote males, although we did not observe any difference in the weights of female *Fam210b^−/−^* mice relative to wild-type controls (Figure 2D). Because of our previous studies suggesting that *Fam210b* is required for erythroid terminal differentiation, we measured splenic weights, as mouse spleens are an extramedullary site of stress erythropoiesis (19). To our surprise, there was no significant difference in the weights (Figure 2E). Peripheral blood smears from adult wild-type and *Fam210b^−/−^* mice did not reveal any gross abnormalities in erythroid cell morphology (Figure 2F).

**Figure 2.**
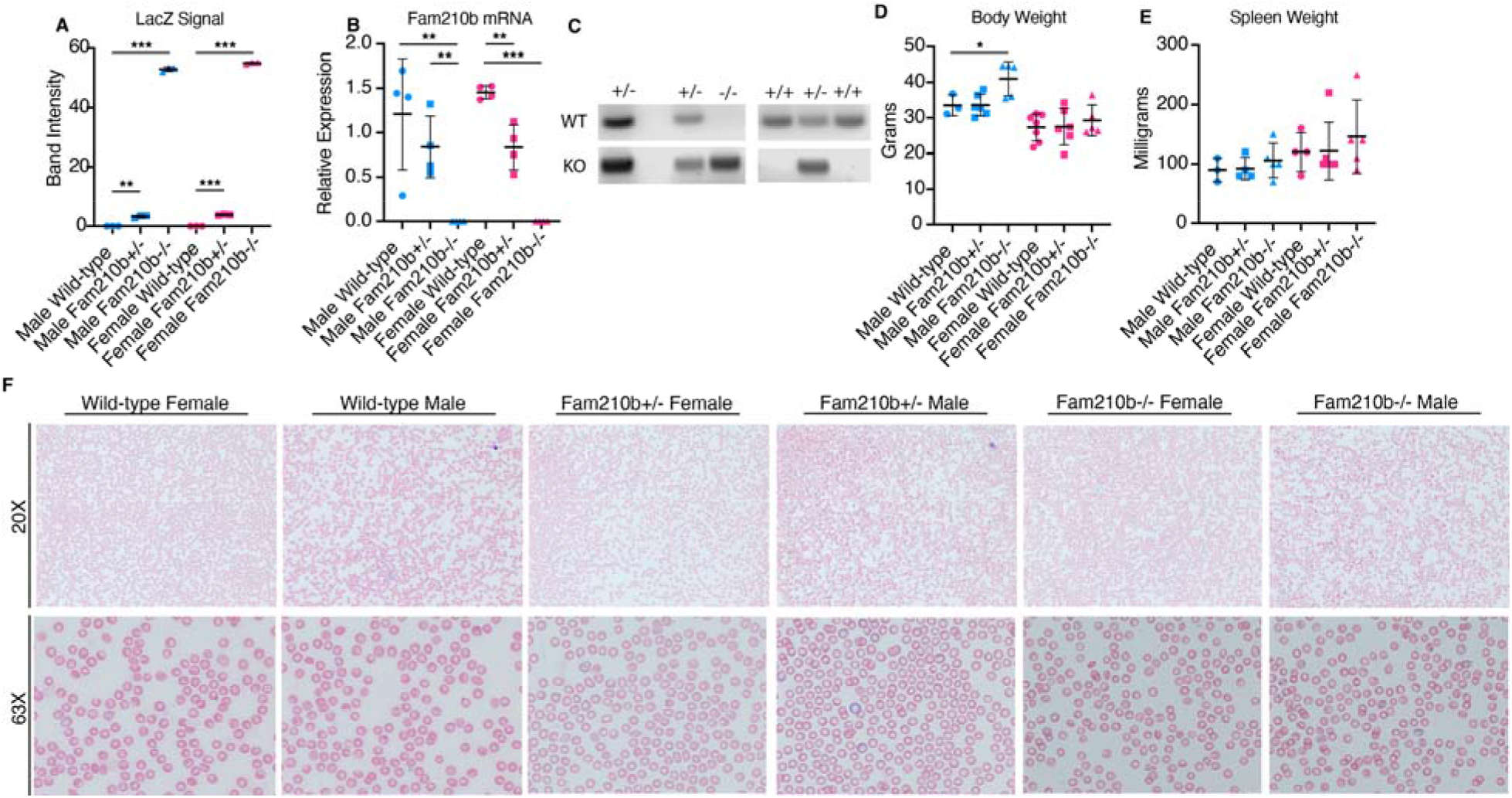
Fam210b deficiency causes a slight increase in male body weight, but has no effect on the gross morphology of the spleen or peripheral blood cells. **A.**, *LacZ* semi-quantitative RTPCR from liver, n=3 per group. **B.**, qRTPCR for *Fam210b* mRNA from liver, n= 4. **C.** PCR from genomic DNA showing excision of the *Fam210b* gene. **D.** Body weight of the mice taken just before dissection, n≥3. **E.**, Weights of mouse spleens, n≥3. **F.** 20X and 63X brightfield images of mouse blood smears stained with May-Grunwald and Giemsa. * P>0.05.

We conducted complete blood count (CBC) analysis to determine if there were any hematological defects in adult *Fam210b^−/−^* mice. We did not observe changes in white blood cell (Figure 3A), lymphocyte (Figure 3B), monocyte (Figure 3C), or neutrophil (Figure 3D) counts. Of interest, there were no significant differences in red cell counts, hemoglobin, hematocrit, mean corpuscular volume, mean cellular hemoglobin or red cell distribution width between WT, *Fam210b^+/−^* and *Fam210b^−/−^* mice of either sex (Figure 3E-K). We did not observe changes in platelet count or platelet volume (Figure 3L-M).

**Figure 3.**
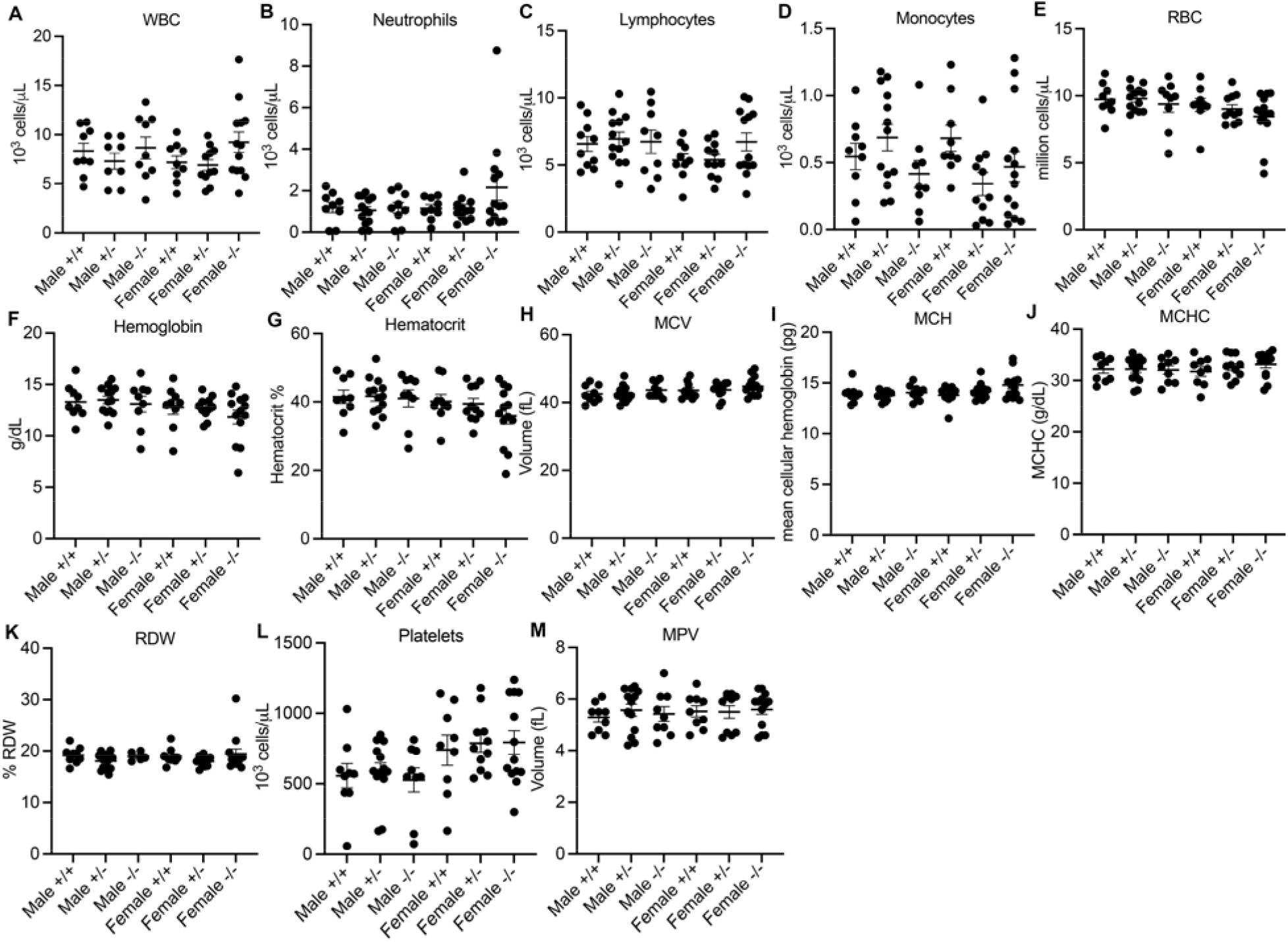
Complete Blood Count of wild-type, Fam210b^+/−^ and Fam210b^−/−^ mice. **A.**, white blood cell (WBC) count, **B.**, neutrophil count, **C.**, lymphocyte count, **D.**, monocyte count, **E.**, red blood cell (RBC) count, **F.**, hemoglobin concentration, **G.**, hematocrit, **H.**, mean corpuscular volume (MCV), **I.**, mean corpuscular hemoglobin (MCH), **J.**, mean corpuscular hemoglobin concentration (MCHC), **K.**, red cell distribution width (RDWs), **L.**, platelet count, **M.**, mean platelet volume (MPV), **N.**, platelet distribution width (PDWs). n≥6 for all values; *, p<0.05.

We then analyzed erythropoietic populations in the bone marrow. Bone marrow smears did not reveal any gross abnormalities between *Fam210b^+/−^* and *Fam210b^−/−^* animals (Supplemental Figure 1). We followed up by quantitating erythroid progenitor populations in the bone marrow (20). Bone marrow erythroid cells were sorted into progenitor populations according to their forward scatter and expression of CD44 and TER119. Populations I-VI corresponded to erythroid cells at progressively mature stages of differentiation (representative FACS plots in Supplemental Figure 1). We did not observe obvious differences in bone marrow erythroid progenitor formation between WT, *Fam210b^+/−^* and *Fam210b^−/−^* mice (Figure 4). Our data indicate that FAM210B is not required for erythroid differentiation or hemoglobinization during steady state.

**Figure 4.**
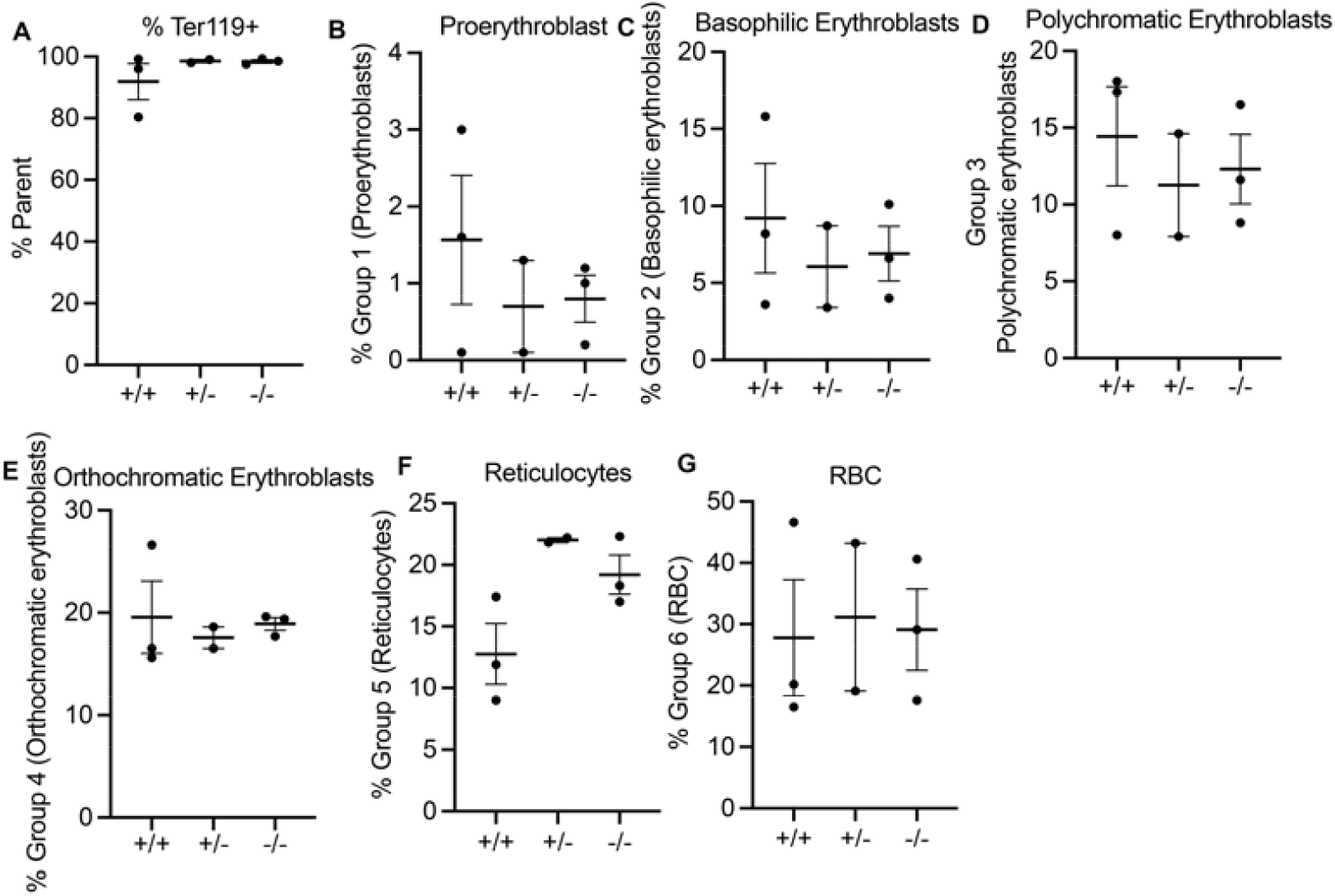
FACS analysis of bone marrow erythroid progenitors in WT, *Fam210b^+/−^* and *Fam210b^−/−^* mice at steady state. We did not observe significant *Fam210b*-dependent differences in bone marrow erythroid progenitors post-phlebotomy; **A.** % Ter119 cells; **B.** Pro-erythroblast; **C.** Basophilic erythroblast; **D.** Polychromatic erythroblast; **E.** Orthochromatic erythroblast; **F.** Reticulocyte and **G.** Mature red cell percentages did not differ between groups.

To determine if *Fam210b^−/−^* animals compensate for erythroid iron deficiency by increasing iron uptake, we measured serum transferrin in WT, *Fam210b^+/−^* and *Fam210b^−/−^* mice. At steady state, transferrin levels were increased in *Fam210b^−/−^* males, but not females (Figure 5A). We did not observe statistically significant differences in steady state serum iron levels between WT and *Fam210b^−/−^* animals (Supplemental Figure 2) indicating the elevated elevated serum transferrin levels may indicate defects in iron sensing in *Fam210b^−/−^* males (21) (Figure 5A).

**Figure 5.**
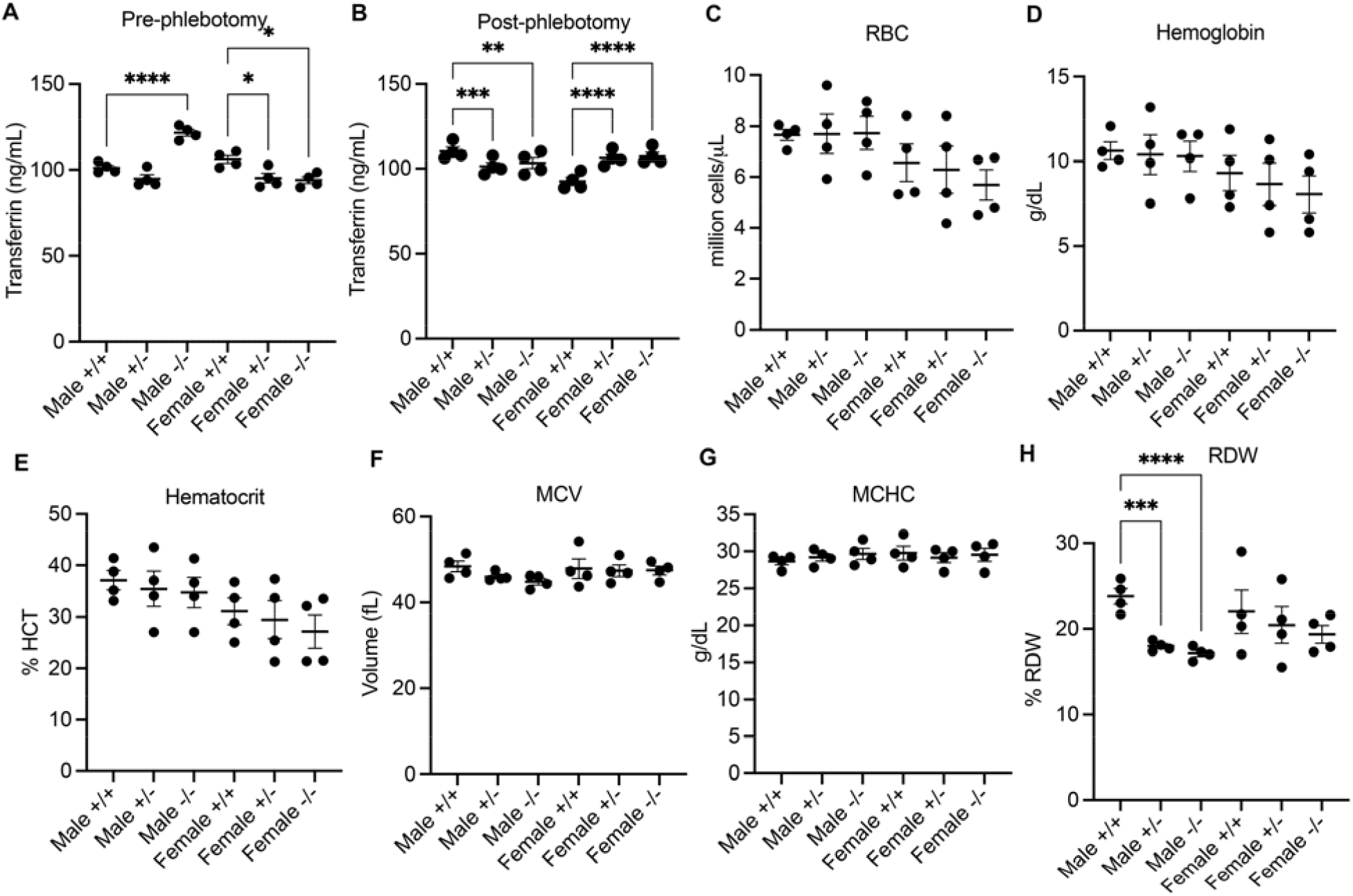
FAM210B is required for iron sensing and response to iron deficiency; however its absence does not cause alterations in red cell production or hemoglobinization during erythropoietic stress. **A.** Pre-phlebotomy transferrin ELISA indicates that male *Fam210b^−/−^* mice have significantly increased transferrin levels, indicating sensing of iron deficiency; this was not observed in females, which have slightly decreased transferrin. **B.** Post phlebotomy, male *Fam210^+/−^* and *Fam210b^−/−^* mice have *decreased* transferrin levels, while females have *increased* transferrin levels. Post-phlebotomy, **C.** RBC; **D.** Hemoglobin; **E.** Hematocrit; **F.** MCV and **G.** MCHC levels do not differ between groups. **H.** Male *Fam210^+/−^* and *Fam210b^−/−^* mice have slightly decreased RDW. *** P = 0.0002; **** P<0.0001

To determine if *Fam210b* was important for erythropoiesis during stress, we carried out serial phlebotomies on WT, *Fam210b^+/−^* and *Fam210b^−/−^* adult mice for iron studies, CBC analyses and FACS analyses of erythroid differentiation. Post phlebotomy, we noticed that transferrin levels of male *Fam210b^+/−^* and *Fam210b^−/−^* were slightly decreased relative to those of wild-type mice, while in the females, *Fam210b^+/−^* and *Fam210b^−/−^* serum transferrin levels were elevated relative to wild-type mice, implicating a sex-specific role in FAM210B function (Figure 5B). As organisms increase transferrin secretion during iron deficiency (21), This was an appropriate response to serum iron concentrations as they were decreased in *Fam210b^+/−^* and *Fam210b^−/−^* females but not males (Supplemental Figure 2). Surprisingly, post-phlebotomy hemoglobin levels and red cell count were not affected by FAM210B status (Figure 5C-H) indicating that decreases in iron may have been offset by increased serum transferrin. Erythroid cell morphology of post-phlebotomy blood did not differ between genotypes or sexes (Supplemental Figure 3). Post phlebotomy CBC indicated significant decreases in neutrophil and monocyte numbers in male *Fam210b^+/−^* and *Fam210b^−/−^* mice (Figure 6).

**Figure 6.**
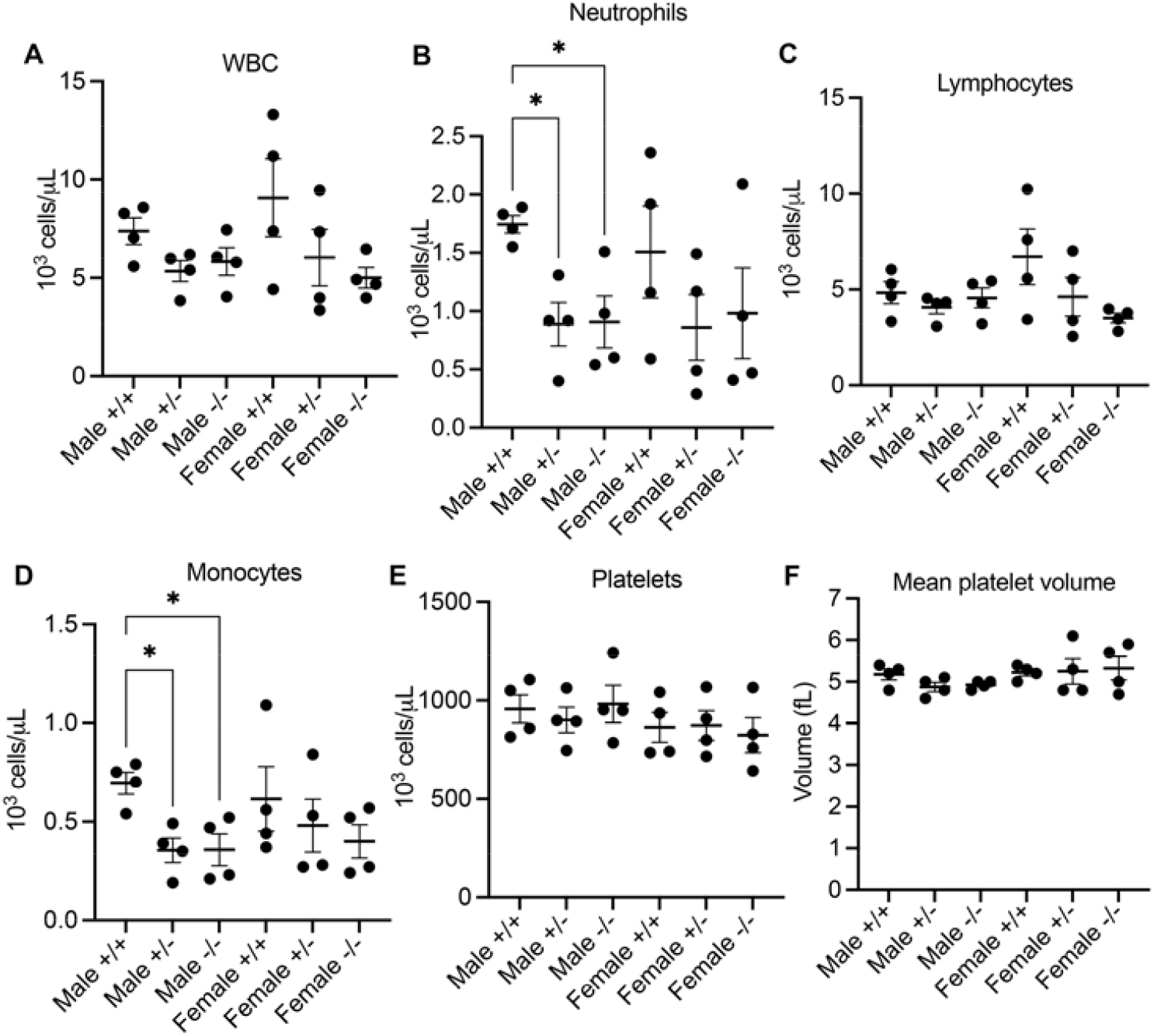
Complete Blood Count of wild-type, Fam210b^+/−^ and Fam210b^−/−^ mice post-phlebotomy. **A.** We did not observe differences in total white cell count, however **B.** neutrophil counts in male *Fam210^+/−^* and *Fam210b^−/−^* mice were decreased post phlebotomy; no differences among female groups. **C.** *Fam210b* was not required for lymphocyte levels, but **D.** Male *Fam210^+/−^* and *Fam210b^−/−^* had decreased monocyte counts, while females were normal. **E.** Platelet counts and **F.** Mean platelet volume was normal in *Fam210^+/−^* and *Fam210b^−/−^* mice. * P<0.05

While the absence of *Fam210b* did not affect bone marrow erythropoiesis during erythropoietic stress (Figure 7), both male and female *Fam210b^−/−^* mice had significantly decreased spleen sizes and weights after erythropoietic stress induced by phlebotomy. This was especially pronounced in the males, where even *Fam210^+/−^* mice had significantly decreased spleen weight relative to body weight (Figure 8A, B). In mice undergoing stress erythropoiesis, short-term bone marrow derived hematopoietic stem cells migrate to the spleen where they are restricted to the erythroid lineage and transiently proliferate (22–24). Hence, this observation was of interest because it indicated that FAM210B plays a role in regulating splenic erythropoiesis during stress. We further interrogated the splenic erythropoietic phenotype by sorting erythroid progenitors with the CD44 and Ter119 markers as described (20). Intriguingly, we observed that male *Fam210b^−/−^* mice had significantly decreased proportions of proerythroblast, polychromatic and orthochromatic erythroblasts relative to their WT counterparts (Figure 8 A-E) but had statistically similar numbers of splenic mature erythrocytes compared to WT males. In contrast, relative proportions of erythroid progenitors appeared unaffected in female *Fam210b^−/−^* mice (Figure 8G). We postulate that male *Fam210b^−/−^* mice accelerate erythroid progenitor development during stress erythropoiesis at the expense of progenitor proliferation accounting for increased proportions of mature erythrocytes by decreasing the progenitor pool yet allowing circulating erythrocyte production to keep pace with wild-type cells. Despite our FACS data (which count a sample of spleen cells), the total number of splenic erythrocytes in both male and female *Fam210b^−/−^* mice significantly lags below wild-type mice as the *Fam210b^−/−^* spleen is about half the weight of a WT spleen (Figure 6). These data are consistent with published observations in *Fam210b* deficient HiDEP erythroid cells, which appeared to have accelerated erythroid differentiation. Our *in vivo* studies suggest that the requirement for FAM210B is specific to splenic derived erythroid cells during erythropoietic stress and is required for expansion of the splenic erythroid compartment during stress. In males, FAM210B is further required for maintenance of the erythroid progenitor pool during stress.

**Figure 7.**
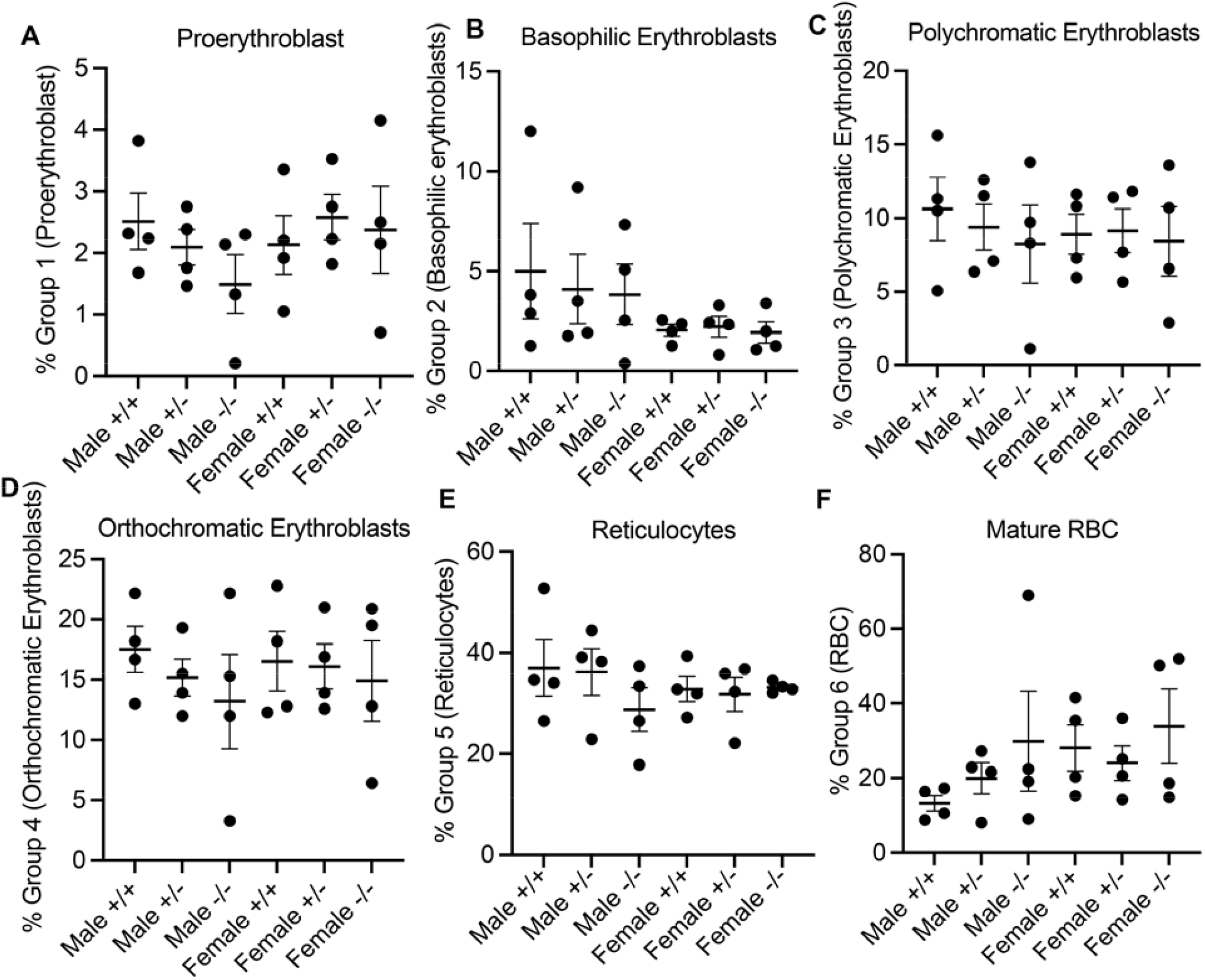
FACS analysis of bone marrow erythroid progenitors in WT, *Fam210b^+/−^* and *Fam210b^−/−^* mice post-phlebotomy. We did not observe significant *Fam210b*-dependent differences in bone marrow erythroid progenitors post-phlebotomy; **A.** Pro-erythroblast; **B.** Basophilic erythroblast; **C.** Polychromatic erythroblast; **D.** Orthochromatic erythroblast; **E.** Reticulocyte and **F.** Mature red cell percentages did not differ between groups.

**Figure 8.**
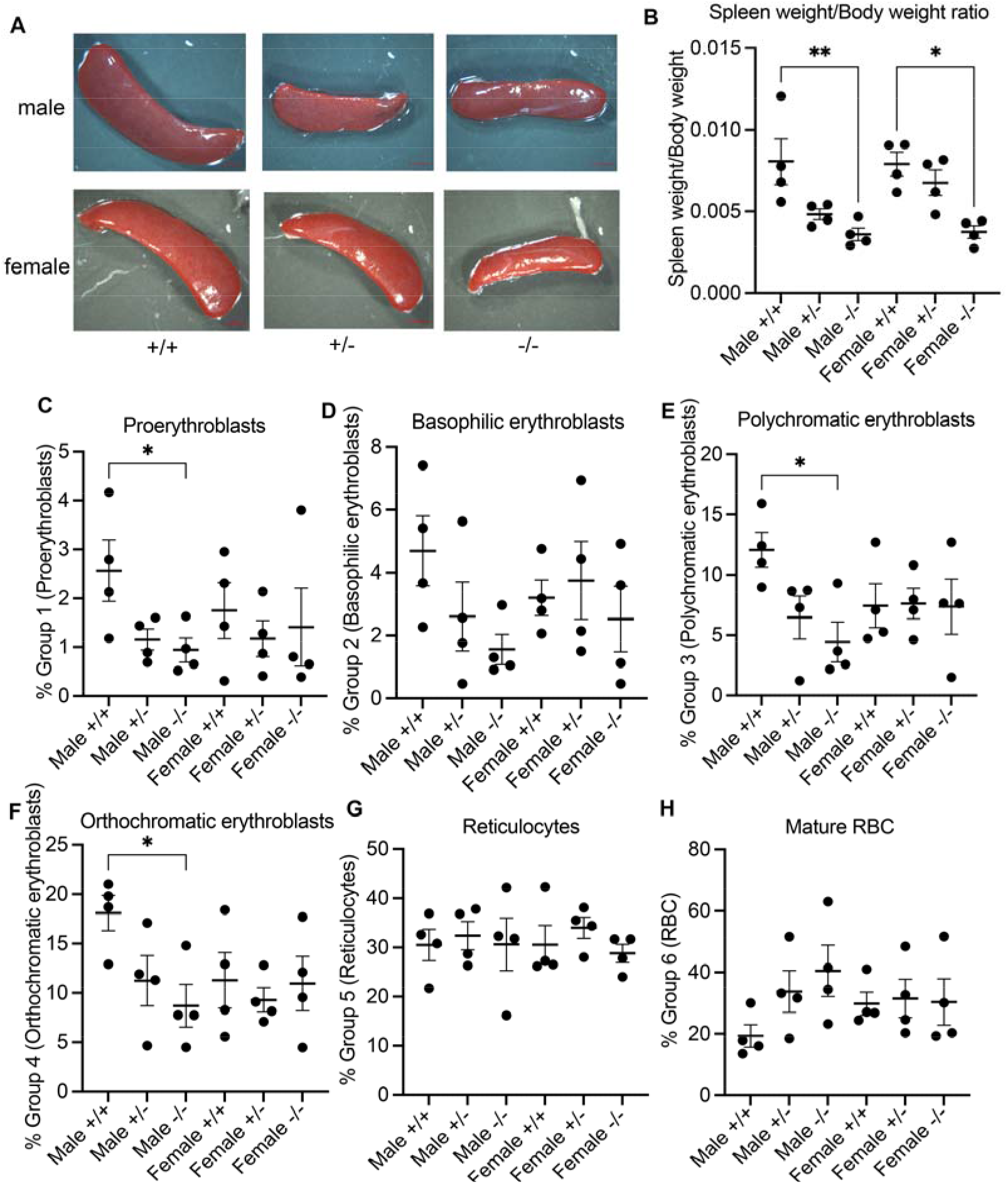
*Fam210b* is required for splenic response to stress erythropoiesis, particularly in male mice. **A.** Photograohs of representative spleens post-phlebotomy (scale bar: 2mM); *Fam210b*^−/−^ spleens are appreciably smaller than wild-type spleens. **B.** When normalized to body weight, *Fam210b*^−/−^ spleens have significantly less mass than wild-type counterparts. FACS analysis of splenic erythroid cells revealed that **C.** Male *Fam210b*^−/−^ mice have significantly decreased percentages of proerythroblasts; females were unaffected. **D.** Basophilic erythroblast populations were unaffected by *Fam210b* deficiency; **E.** Male *Fam210b*^−/−^ mice have significantly decreased percentages of polychromatic erythroblasts but females were unaffected; **F.** Male *Fam210b*^−/−^ mice have significantly decreased percentages of polychromatic erythroblasts but females were unaffected. All groups had similar numbers of **G.** splenic reticulocytes and **H.** red cells. * P<0.05; **P<0.01

## Discussion

This manuscript describes the first study of FAM210B’s role in adult erythropoiesis, identifying it as the first sex-specific regulator of stress erythropoiesis in vertebrates. Based on previous work, we expected that *Fam210b* knockout phenotype would be pre-embryonically lethal due to its requirement for fetal liver erythroid hemoglobinization and differentiation in primary cell culture (5). We were therefore surprised that *Fam210b* was dispensable for reproduction and viability in mice. The survival of *Fam210b^−/−^* mice to adulthood suggested that erythropoietic defects caused by *Fam210b* deficiency were compensated by genetic or environmental factors. We demonstrated that holotransferrin rescued hemoglobinization of *Fam210b^−/−^* MEL cells even when cells were grown in 10% FBS (Figure 1). These data are consistent with our previous results demonstrating that supplemental iron restores the erythroid differentiation and hemoglobinization defects in *Fam210b^−/−^* erythroid cells that were cultured in 10% FBS, supporting the idea that one can bypass the need for *Fam210b* in embryos by supplying bioavailable iron.

In vertebrates, erythroid cells predominantly obtain iron from holo-transferrin in niches known as erythroblastic/erythromyeloblastic islands where developing erythroid cells remain in close spatial proximity to a central macrophage (25,26). The central macrophage of the erythroblastic island is a major source of iron for differentiating erythroid cells (27–29), expressing high levels of iron transport proteins such as transferrin, ferroportin and ferritin (21) which creates localized regions of highly concentrated bioavailable iron. Therefore, it is probable that high local concentrations of iron are able to circumvent hemoglobinization defects in *Fam210b^−/−^* erythroid cells. Further, bovine transferrin has very low affinity to murine transferrin receptor, creating iron deficient conditions under the standard 10% FBS cell culture condition by default compared to holotransferrin concentrations in vertebrates (30,31). Serum iron concentrations in males are not altered by *Fam210b* deficiency (Supplemental Figure 2), but *Fam210b* deficiency appears to cause alterations in iron sensing even at steady state. Male *Fam210b^−/−^* mice have significantly higher serum transferrin levels than wild-type mice, suggesting that they sense iron deficiency; these increased transferrin levels may compensate for any decreased iron supply to erythroid cells. In contrast, female *Fam210b^−/−^* mice have slightly decreased serum transferrin levels, indicating they do not similarly sense iron deficiency

The hypoxic environment of the fetal liver and bone marrow increases intracellular iron uptake relative to normoxic conditions (such as in standard tissue culture) via the upregulation of genes such as *Dmt1* and *TfR1* (32–36). In contrast, zebrafish embryos grow under normoxic conditions where tissue oxygenation initially occurs by passive diffusion prior to blood formation (37), and their erythroid cells do not form erythroblastic islands. Zebrafish obtain iron from their yolk stores, before they begin eating at 5-6 dpf. In contrast, mammalian embryos obtain their iron from maternal circulation (38). These factors likely underlie why *fam210b-* deficient zebrafish erythrocytes appeared severely anemic, while *Fam210b^−/−^* fetuses were able to produce sufficient numbers of erythrocytes for viability to the point of birth.

We tested the hypothesis that FAM210B primarily functions to adapt to iron deficiency or erythropoietic stress. Mice that were subjected to serial phlebotomy which depletes their body iron stores, had defects in their stress erythropoiesis response (Figures 5, 6, 8). Although their erythroid indices post phlebotomy were normal, there were sex-dependent differences in their iron sensing. Female *Fam210^+/−^* and *Fam210b^−/−^* mice had increased serum transferrin consistent with decreased iron (Supplemental Figure 2), while the males had decreased serum transferrin relative to wild-type mice (Figure 5). CBC analyses of their peripheral blood indicated decreased neutrophil and monocyte counts in male *Fam210^+/−^* and *Fam210b^−/−^* mice post phlebotomy suggesting *Fam210b*-dependent mechanisms for maintaining neutrophil and monocyte populations in males during stress.

A key finding in this manuscript is the requirement for *Fam210b* in the erythropoietic stress response. In situations of increased erythropoietic demand, splenic erythropoiesis plays a dominant role in responding to stress causing a dramatic increase in spleen size due to vastly increased rates of erythropoiesis (39). Both male and female *Fam210b^−/−^* mice had spleen weights approximately half that of wild-type mice, indicating their spleens did not expand as they normally would in response to stress (Figure 8). FACS quantitation of splenic erythroid cells indicated that proportionally, female *Fam210b^−/−^* splenic erythroblasts did not have developmental defects, although total cellular numbers were decreased compared to wild-types because of their decreased spleen size. Intriguingly, male *Fam210b^−/−^* spleens had significantly decreased percentages of proerythroblasts, polychromatic and orthochromatic erythroblasts and normal percentages of reticulocytes and mature red cells, suggesting that they responded to stress by accelerating through the terminal steps of erythropoiesis, consistent with observations by Suzuki *et al.* (15). Interestingly, the decreased splenic erythroid cell production was sufficient to maintain wild-type levels of red cells and hemoglobin (Figure 5) over the relatively short assay period, though extended periods of iron deficiency may have caused more severe *Fam210b^−/−^* specific phenotypes. Alternatively, egress of erythrocytes from the spleen may have been accelerated as a compensatory mechanism. We speculate that the defect in splenic erythropoiesis may have contributed to the lack of a severe serum iron defect in *Fam210b^−/−^* mice (Supplemental Figure 2) as a massive proliferation of splenic erythrocytes would have consumed a large quantity of iron.

We did not observe any obvious developmental or functional defects in *Fam210b^−/−^* mice. Male *Fam210b^−/−^* were heavier than their wild-type and female counterparts, which warrants closer investigation of body composition and cellular oxygen consumption. Suzuki et al. reported that FAM210B plays a role in ATP generation in erythroid cells via metabolic reprogramming which is of interest moving forward. Of note, substrate concentration in anatomical and developmental niches (such as iron in the erythroblastic island) can sometimes mask defects in metabolic genes. For example, ATPIF1/IF1, initially observed to be important for erythropoiesis in zebrafish embryos (40), was shown to be dispensable for development and reproduction in mice (41). Another hypothesis is that there may be other uncharacterized compensatory mechanisms that exist in mammals that have not been fully appreciated.

Our studies indicate a male-specific role for FAM210B in splenic erythropoiesis and production of neutrophils and monocytes during stress. Hormonal control of hematopoiesis has long been observed in vertebrate organisms (42–44) but the mechanisms are poorly understood. To determine if *Fam210b* could be regulated by androgens, we performed an analysis of human androgen receptor (AR) ChIPseq data from Pomerantz et al (45). AR was recruited to multiple sites in the *FAM210B* promoter in both normal and cancerous prostate (Supplemental Figure 4), suggesting that testosterone may regulate *Fam210b* transcription. Our data indicates that *Fam210b* gene regulation may be one mechanism by which testosterone regulates stress erythropoiesis and iron homeostasis.

FAM210B’s role in cellular homeostasis is highly complex. Our data indicates that careful control of cell culture conditions is essential for carrying out and interpreting studies on nutrient metabolism *ex vivo*. Further, our studies underscore the importance of carrying out studies in males and females to uncover underlying differences in physiology. We report that FAM210B is dispensable for steady state erythropoiesis and iron uptake, but regulates stress erythropoiesis and iron sensing, particularly in males. Our studies report the first known sex-specific regulator of erythropoiesis in vertebrates. The ability of mice to function in the absence of FAM210B also raises the possibility that human FAM210B mutations may function as modifiers of disease severity in patient disorders of iron metabolism or erythropoiesis as these organisms are viable and fertile, and can pass on such mutations to their progeny.

## Methods and materials

### Cell culture

Murine erythroleukemia (MEL) cells were a gift of Dr. Arthur Skoultchi at the Albert Einstein College of Medicine. The *Fam210b^−/−^* MEL cells were previously generated as described (5). Cells were cultured in DMEM with high glucose, supplemented with 1% glutamine and 10% FBS unless otherwise indicated.

### Heme staining

*o-*Dianisidine staining was performed as previously described (46). Cells were stained in the dark for 10 minutes at room temperature unless otherwise indicated. Imaging was conducted using a Zeiss Axio Imager.A2 and ZEN Blue software (version 3.1). Heme staining was quantitated using ImageJ.

### Cell size analysis

2.5*10^5^ MEL cells were mounted onto a slide and imaged with a Zeiss Axio Imager.A2 and ZEN Blue software (version 3.1). Cell size was calculated in ImageJ by measuring the area of individual cells.

### Mice

*C57BL6N/6N-^Atm1Brd^ Fam210b^tm1b(KOMP)/Mbp^*/MbpMmucd (*Fam210b^−/−^*) whole body knockout mice were generated by UC Davis and purchased from MMRRC. *Fam210b* was genotyped by Transnetyx using primers flanking exon 2 with the following sequences: forward: CTCTGGGTCAGCGAGACAAC, reverse: AGGACTGAGACCCAGGCAAT. Mice were maintained at the University of Delaware and University of Pittsburgh Animal Facilities and according to institutional guidelines. Mice used in these studies were between 3 to 6 months of age. The WT *Fam210b* allele was amplified by the following primers: forward: TGGAAAAGCTCTCACAGACCTACCC; reverse: CTGCCATTTTGGACTGCACCAGG. The *Fam210b^−/−^* allele was amplified by the following primers: forward: GCTACCATTACCAGTTGGTCTGGTGTC; reverse: CTGCCATTTTGGACTGCACCAGG

### RNA Isolation and cDNA generation from mice

RNA was isolated from mouse tissue using the RNAeasy kit (Qiagen). cDNA was then immediately generated using the high-capacity cDNA reverse transcription kit (Applied Biosystems).

### RT-PCR analysis

The following Taqman probes from Applied Biosciences were used for qRT-PCR analysis: *Actb-* VIC (Mm02619580_g1) and *fam210b-*FAM (Mm01248184_m1).

RT-PCR for *lacz* was then conducted using the HOTSTAR protocol (Thermo) with forward primer sequence: 5’ CTCCACAAGGATAACAGTTGA and reverse primer sequence: 5’ ACTACCATCAATCCGGTAGGT. Electrophoresis of PCR products were done on a 1% agarose gel and analysis was conducted on a LI-COR Odyssey Fc and Image Studio software (version 5.2; LI-COR Biosciences).

### Blood and bone marrow extractions

Blood was extracted by doing a cardiac puncture using EDTA coated needles and syringes. At least 100uL of blood was gathered and slowly added to an EDTA coated blood collection tube (K3 EDTA micro500; SAI Infusion Technologies). The blood was kept at 4°C and was analyzed and processed further within 2 hours of extraction.

Bone marrow was extracted from the long bones of the mice. In brief, tissue surrounding the bone was removed with sterile scissors and scalpels. The bone caps were then cut off using scissors and the marrow was removed by gently passing ice-cold 1mL 1X PBS with 0.5% BSA and 2mM EDTA through the bone cavity and into a microcentrifuge tube using a 1mL EDTA coated syringe capped with a 27G EDTA coated needle. The bone marrow was kept at 4°C and was analyzed and processed further withing 6 hours of extraction.

### Blood and bone marrow smears

Thin blood and bone marrow smears were made by placing a 10uL of blood or isolated bone marrow solution (see above) on a charged slide near the frosted end. Another slide was brought up to the drop at a 30-45° angle and the drop was allowed to spread along the contact line between the slides. The angled slide was then quickly dragged toward the unfrosted end. The blood and bone marrow were allowed to dry overnight at room temperature and then stained using May-Grunwald solution (Sigma-Aldrich) for 5 minutes followed by 3 washes in 40mM Tris buffer (pH=7.2) and then 1:20 Giemsa solution (Sigma-Aldrich). The blood and bone marrow were then imaged on a Zeiss Axio Imager.A2 and Zen Blue software (version 3.1).

### Complete blood count

A complete blood count was conducted using an Abaxis Vetscan HM5 Hematology Analyzer (version 2.3) according to the manufacturer’s protocol.

### Erythroid progenitor flow cytometry

Erythroid progenitor cells were sorted as described previously (20). In brief, single-cell bone marrow suspensions (see above) were centrifuged at 4°C, 300g for 10 minutes. For spleen samples, we ∼20mg of spleen in 1.5mL tubes and kept samples on ice. 100uL of ice cold 1X PBS with 1% BSA was added to the spleen and then dissociated using a 1.5mL pestle. 900uL of ice cold 1X PBS with 1% BSA was added to the spleen dissociate. The cells were then counted on a hemacytometer and sorted (20).

CD45 was depleted by resuspending 10^7^ cells with 100uL of a 10% CD45 MicroBead Slurry (MiltenyiBiotec) in 1X PBS with 0.5% BSA and 2mM EDTA. CD45 depleted bone marrow suspensions were blocked with rat anti-mouse CD16/CD32 (BD Bioscience) and stained using BD Bioscience antibodies: TER-119-FITC, CD44-APC, CD45-APC-Cy7, CD11b-APC-Cy7, GR1-APC-Cy7, and 7AAD. The bone marrow cells obtained after bead depletion were imaged in Supplemental Figure 1.

For flow cytometry, stained bone marrow suspensions removed dead cells by gating out 7AAD positive cells (Cy5). From this population, erythroid progenitor cells were selected by gating out APC-Cy7 positive immune cells cells (CD45, GR1, CD11b) and cells with high internal complexity by SSC-A. The remaining cells were then selected for TER-119 expression (FITC) and then this population was sorted by CD44 (APC) and TER119 (FITC). Erythroid progenitor groups I-VI were then selected and sorted from this population. For imaging, cells were sorted directly onto a slide and allowed to dry overnight. The cells were then stained using May-Grunwald solution and Giemsa as described above and imaged on a Zeiss Axio Imager.A1 and Zen Blue Software (version 3.1).

### Cell cycle analysis of MEL cells

Cells were harvested and then washed three times with PBS followed by fixation with ice-cold 70% ethanol for 30 minutes on ice. The cells were then rehydrated and washed 3 times with 1X PBS. The RNA was digested using 100ug/mL RNase and DNA was stained using 40ug/mL propidium iodide. The cells were then analyzed for cell cycle phase by flow cytometry.

### Phlebotomy and Cardiac Puncture

∼500uL of blood was collected once per day over three days by submandibular bleed. The blood was collected into an EDTA coated tube for CBC analysis and a 1.5mL Eppendorf tube for serum iron analysis. Bleeding was stopped either naturally or by pressure with a sterile cloth. Cardiac puncture to collect post-phlebotomy blood was performed 3 days after the last submandibular bleed. The mice were then euthanized in accordance with IACUC protocols.

### Serum Iron Analysis

SEKURE Chemistry Iron-SL Assay (SEKURE 157-30) was performed on serum from clotted blood. After collection, the blood was clotting by incubating it at room temperature for 30 min. The blood was then centrifuged 4500rpm for 5 minutes and the serum was aspirated. The serum was centrifuged again, and the aspirate was collected. The serum was then snap frozen in liquid nitrogen and stored at −80C until needed. Serum iron was then analyzed according to the manufacturer’s protocol.

### Serum Transferrin ELISA

Serum was collected from clotted blood as noted above. The serum was then analyzed using a mouse Transferrin ELISA kit and in accordance with the manufacturer’s protocol (Alpha Diagnostic International 6390).

## Statistical analysis

Statistical significance was carried out using Prism and assessed by one way ANOVA with Tukey’s correction for multiple comparisons.

## Acknowledgements

We thank the late Richard West for assistance with flow sorting and use of the CBC counter and Dr. Gwen Talham for extensive, invaluable assistance with mouse work during the 2020 COVID-19 lockdown. We thank the University of Pittsburgh flow cytometry and mouse cores for technical assistance. We are grateful to Drs. Molly Sutherland and Samit Ghosh for helpful comments, and members of the Yien and Tejero labs for feedback. This work is supported in part by PHS grants P01HL032262 (Project 5), R35GM133560 and R01 DK134783 (YYY), R35GM137979 (ANS) and T32HL149648 (MP; PI Novelli), the Cooley’s Anemia Foundation (YYY), and P3VHB funding (JT).

