## Supplemental data for "FAM210B Regulates Iron Homeostasis and Sex-Specific Responses in Stress Erythropoiesis"

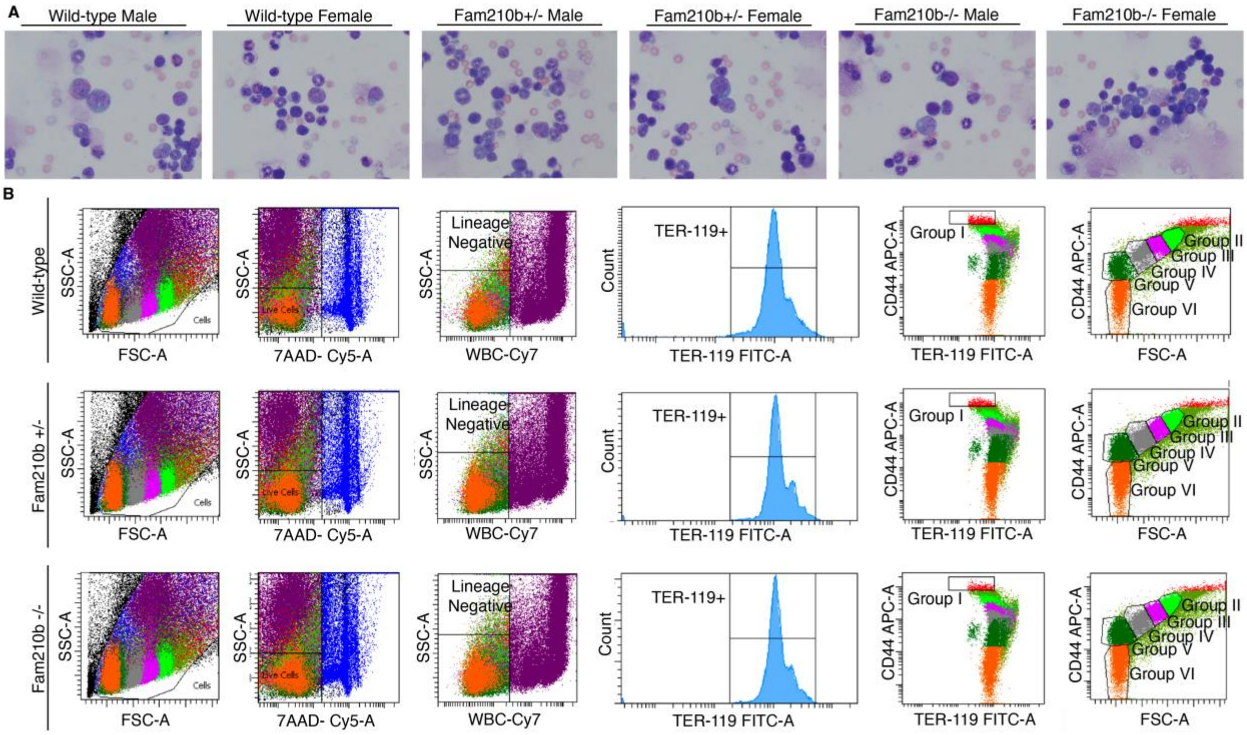


**Supplemental Figure 1. Fam210b deficiency does not affect bone marrow gross morphology or erythroid progenitor formation. A.**, 20x brightfield images of Grunwald/Giemsa stained mouse bone marrow cells. **B.**, Flow cytometry gating for quantification of erythroid progenitor populations; Group I: Pro-E, Group 2: Baso-E, Group III: Poly-E, Group IV: Ortho-E, Group V: Reticulocytes, Group VI: RBC.

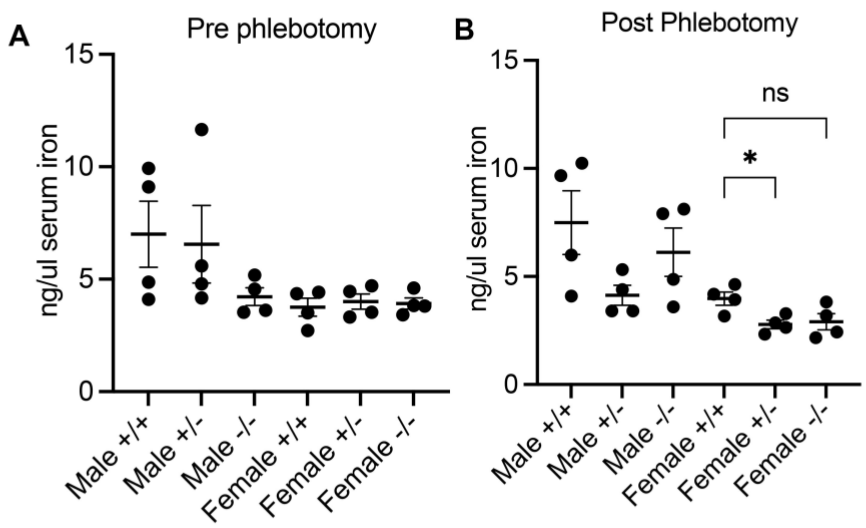


**Supplemental Figure 2. Serum iron concentration in *Fam210b*$^{+/+}$, *Fam210b*$^{+/-}$, *Fam210b*$^{-/-}$ mice.** **(A)** Pre-phlebotomy, serum iron across genotypes did not differ from one another. **(B)** Post-phlebotomy, female *Fam210b*$^{+/-}$ mice had significantly decreased serum iron relative to wild-type; *Fam210b*$^{-/-}$ mice trended to having decreased serum iron (NS: $p = 0.075$). * $p<0.05$

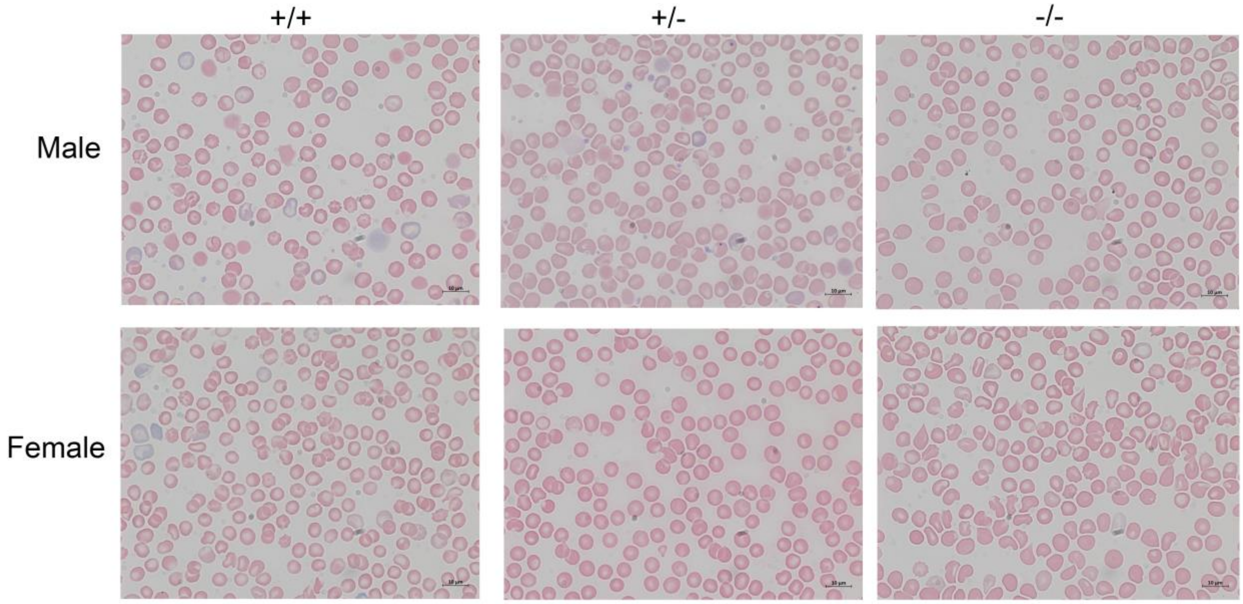


**Supplemental Figure 3. Blood smears from *Fam210b*$^{+/+}$, *Fam210b*$^{+/-}$, *Fam210b*$^{-/-}$ mice post-phlebotomy.** There is no difference in red cell morphology between the genotypes post-phlebotomy. Images taken at 63x magnification; scale bar = 10µm.

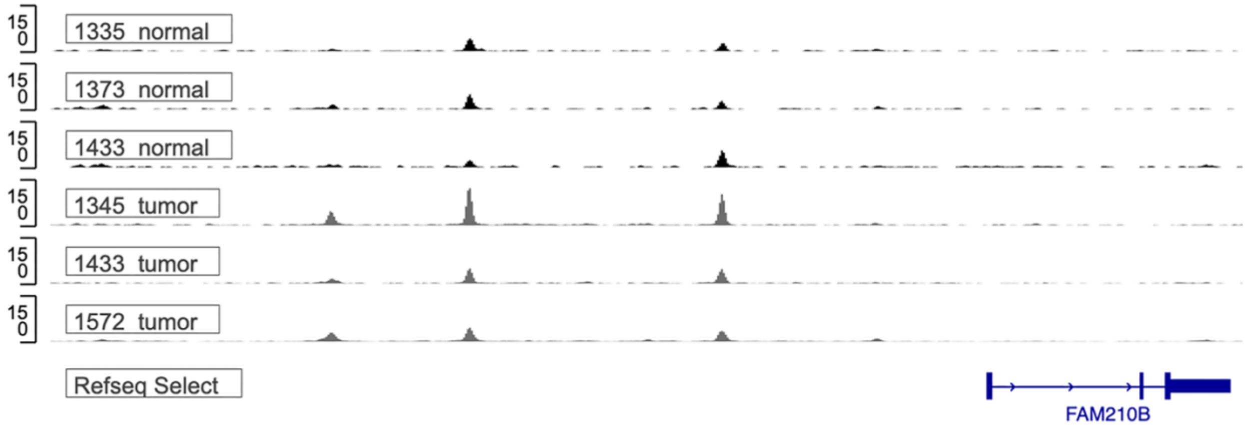


**Supplemental Figure 4. Recruitment of AR to *FAM210B* promoter in human prostate.** ChIPseq tracks from three normal and three cancerous prostate samples show AR binding to multiple sites upstream of *FAM210B*.
